# ARGs-OSP: online searching platform for antibiotic resistance genes distribution in metagenomic database and bacterial whole genome database

**DOI:** 10.1101/337675

**Authors:** An Ni Zhang, Chen-Ju Hou, Li-Guan Li, Tong Zhang

## Abstract

**Background:** The antibiotic resistant genes (ARGs) have been emerging as one of the top global issue s in both medical and environmental fields. The metagenomic analysis has been widely adopted in ARG-related studies, revealing a universal presence of ARGs in diverse environments from medical settings to natural habitats, even in drinking water and ancient permafrost. With the tremendous resources of accessible metagenomic datasets, it would be feasible and beneficial to construct a global profile of antibiotic resistome as a guidance of its phylogenetic and ecological distribution. And such information should be shared by an open webpage to avoid the unnecessary repeat of data processing and the bias caused by incompatible search method.

**Results:** Two dataset collections, the Whole Genome Database (WGD, 54,718 complete and draft bacterial genomes) and the Metagenomic Database (MGD, 854 metagenomic datasets of 7 eco-types), were downloaded and analyzed using a standard method of ARG online analysis platform (ARGs-OAP v1.0). The representativeness of WGD and MGD was evaluated to have a comprehensive coverage of ARGs in bacterial genomes and metagenomes. Besides, an ARGs online searching platform (ARGs-OSP, http://args-osp.herokuapp.com/) was developed in this study to make the data accessible to other researchers via the search and download functionality. Finally, flexible usage of the ARGs-OAP was demonstrated by evaluating the co-occurrence of class 1 integrases and total ARGs across different environments.

**Conclusions:** The ARGs-OSP is presented in this study as the valuable sources and references for future studies with versatile research interests, meanwhile avoiding unnecessary re-computations and re-analysis.

## Background

Due to the intensive usage of the human and veterinary antibiotics, the antibiotic resistant genes (ARGs) are emerging in almost all environments as one of the top global concerns. Recently, through high-throughput sequencing and metagenomic analysis, the ARG profiles have been investigated in diverse habitats, especially the anthropogenic environments of human microbiome [1–6], animal microbiome [7–10] and WWTPs [11–13]. ARGs, especially clinically relevant ones [14–18], have been spreading from the anthropogenic habitats to the natural eco-systems [19], mainly contributed by WWTPs, pharmaceutical manufacturing plants, hospitals, and husbandry facilities [13, 20–23]. Universal presence of various ARGs has been revealed in all kinds of natural ecosystems by many metagenomic studies investigating the samples of sediment [23–25], soil [26–28], surface water [29–31], marine water [27] and even ancient permafrost [32–34]. The identification of ARGs in drinking water [35] and human food [36, 37] further reveals the potential of their direct exposure to human health. With the growing attention to the environmental issue of antibiotic resistance dissemination all over the world, ARG-related studies have gained momentum, and are covering all kinds of habitats. All these metagenomic datasets are valuable and accessible sources to construct a global profile of antibiotic resistome.

Despite the recent increase in aforementioned metagenomic datasets, the approaches of the identification and annotation of ARGs varied in different researches regarding the searching methods, searching criteria, the reference databases and the units in the quantification. This makes direct inter-sample comparison infeasible. For example, the ARG profiles were evaluated to have significant difference of up to 5-20 fold [38] when using the domain-based searching method of Hidden Markov Model (HMM) [39] against the similarity-based searching approaches of BLAST [40], USEARCH [41] and DIAMOND [42]. Another major obstacle for the parallel comparison was the use of different reference databases. Even for the top two highly-cited ARG databases, the Comprehensive Antibiotic Resistance Database (CARD; http://arpcard.mcmaster.ca) [43] and Antibiotic Resistant Database (ARDB) [44], bias could be raised during comparison because they only share a small portion of reference sequences [45]. Recently, a widely-applied and well-curated ARG database named the Structured ARG reference database (SARG; http://smile.hku.hk/SARGs) [38, 45], was constructed by integrating both the CARD and ARDB, which was selected as a standard database in this study. The metagenomic datasets from diverse habitats should be re-analyzed using a standardized pipeline for ARG identification and annotation, from which the ecological distribution of ARGs could be comprehensively drawn.

Although metagenomic analysis could uncover the distribution and abundance of ARGs in a habitat, the information it can provide is limited. Further information describing the host, the mobility and the gene arrangement of ARGs is critical and necessary to investigate the frontier scientific questions about the origin, the evolution, the spreading and the co-selection of ARGs in different environments. This problem can be partially alleviated through the use of assembled metagenomes and nanopore sequencing techniques, but such studies are still limited to only a few environmental samples [35]. On the contrary, such details can be provided with precision and certainty through analyzing the collection of bacterial whole genomes. It was demonstrated by some previous researches [46, 47] via mining the co-occurrence patterns of ARGs and metal resistant genes (MGEs) in the collection of bacterial complete genomes. To construct a global profile of ARGs, the integration of whole genomes and metagenomes is a promising attempt.

Many public databases publish the information in a well-organized way for the convenient search and download by the user ends, such as the IMG/VR (https://img.jgi.doe.gov/vr/) [48]. However, to the best of our knowledge, such application was not introduced to the field of ARG research, especially to provide information on the phylogenetic and ecological distribution of antimicrobial resistance. Moreover, the processing of the big datasets of whole genomes and metagenomes is time- and resource-consuming, and is unnecessary repeated by individual researchers all over the world. In this study, a global profile of ARGs is constructed by a standard pipeline and is presented in the form of an online searching platform for ARGs (ARGs-OSP, http://args-osp.herokuapp.com/), serving as a valuable resource for future studies.

## Data and Webpage Description

To construct a global ARG profile covering the information of their phylogenetic and ecological distribution, the ARGs were identified and quantified by searching two collections of bacterial whole genomes and metagenomes using a standard pipeline, the ARGs online analysis platform (ARGs-OAP v1.0) [45]. The occurrence and abundance of ARGs were summarized and organized with the metadata information into mothertables, which were published on the ARGs-OSP (http://args-osp.herokuapp.com/). On ARGs-OSP, search and download functionality was designed for users to retrieve the occurrence of ARGs in different taxonomy and the abundance of ARGs in different habitats. The availability and convenience of this platform could meet the requirements of versatile research interests, such as: the current host range of some specific resistant genes on the bacterial phylogenetic tree, the dissemination of some specific resistant genes in both the natural and anthropogenic habitats, the antibiotic resistome currently detected in some specific taxa or specific environments, and the comparison of the ARG profiles of a local sample to the global profile. Through data sharing, ARGs-OSP is expected to motivate and facilitate future studies into mining new information and knowledge from the combined data, without making repeated efforts in dataset processing.

## Methods

### Two collection of datasets

The Whole Genome Database (WGD) containing 54,718 bacterial genomes (7,770 complete genomes and 46,948 draft genomes with medium and high quality of more than 50% completeness [49]) was downloaded from the NCBI genome database [50] (ftp://ftp.ncbi.nlm.nih.gov/genomes/genbank/bacteria, ftp://ftp.ncbi.nlm.nih.gov/genomes/GENOME_REPORTS/prokaryotes.txt) (on July 8, 2017), as summarized in Table S1. The potential pathogenicity of bacterial genomes was obtained by matching their taxonomic information with a published database covering the taxonomy of currently recognized human bacterial pathogens [51] (Table S1). If either one of the taxonomic annotations of genus, species or strain level was matched to the human pathogen list, the genome was labeled as a potential human pathogen [10, 52].

The Metagenome Database (MGD) totaling 854 metagenomic datasets were downloaded from NCBI SRA database [50] and MG-RAST[53], which were all generated through Illumina shotgun sequencing. The habitat information of the metadata of all samples was manually organized to categorize them into totally 25 eco-subtypes of 7 eco-types by integrating the guidance of previous studies [54, 55], covering both the natural environments (water, sediment, soil, and permafrost) and the anthropogenic environments (WWTPs, animal feces, and human feces). Thus, this collection of MGD is expected to represent a wide and comprehensive ecological diversity. The quality control of raw reads was conducted with Fastx-Toolkit (http://hannonlab.cshl.edu/fastx_toolkit) with the minimum quality score of Q20 within at least 90% of bases, resulting in in the total number of clean reads varied from 0.1 million to 91 million. All raw reads were trimmed equally to the length of 100bp to allow more accurate inter-sample comparison.

### Identification and annotation of the ARGs and intl1 gene

All the coding sequences (CDS) of WGD were extracted from the genbank files to be searched against SARG [45] and the class 1 integrases *(intI1)* database (manuscript under review). Those CDS meeting the criteria of e-value 1e-5, 90% amino acid (aa) identity over 80% aa hit-ratio against the SARG, and e-value 1e-3, 80% aa identity over 50% aa hit-ratio against the *intI1* database, were annotated as the ARGs and *intI1* genes, respectively. The ARGs and *intI1* genes in MGD were also investigated by these two databases. An effective and time-saving searching process was conducted by two-step sequence-based methods, first through usearch v8.0.1623_i86linux64 [41] and followed by BLASTX 2.2.28+ [40]. The abundance of the ARGs and *intI1* genes was calculated and transformed to different units of ppm, copy per 16S and copy per cell. Furthermore, the abundance of each unit was specifically calculated under different combination of searching criteria (e-value, identity and hit-length). ARGs-OSP provides 60 combinations of searching criteria for each abundance unit, that is, totally 180 mothertables for more flexible usage by future studies. The cutoff used in previous studies [7, 45] (manuscript under review) for metagenomes was e-value 1e-7, 80% aa identity over 75% aa hit-length of the SARG and the *intI1* database, which was adopted in this study as a standard cutoff for further analysis. Also, the standard abundance unit for the ARGs and *intI1* genes in MGD was set as copy per cell, which was comparable to copy per genome in WGD.

### Mothertable analysis and visualization

The mothertables were organized by combining the ARG profile with the phylogenetic and ecological information, via self-written Python 2.7.6 and R scripts using R 3.3.2[56] (packages’dplyr’,’ggalt’,’ggthemes’, ‘ggplot2’, and ‘plyr’) and Python 2.7 (https://www.python.org/). The rarefaction curves[57] of WGD (based on each genome) and MGD (based on each raw read) were conducted by randomly subsampling the genomes or raw reads without replacement [58, 59], that is, each genome or raw read was sampled only once. The step of the raw read number for the entire MGD and for each eco-type was set at the total number of raw reads (for each dataset) divided by 10,000, which ensured that each rarefaction curve was plotted with 10,000 points. All the networks were visualized with Cytoscape 3.3.0 [60] in *Tree* or *Hierarchic* layout or with R 3.3.2[56] (package ‘ggplot2’).

## Results and Discussions

### The representativeness and coverage of WGD and MGD

In WGD, 54,718 bacterial genomes were downloaded from the NCBI genome database in total [50], covering 32 bacterial phyla, 162 classes, 299 orders, 643 families, 1,986 genera, and 3,654 species, without counting the unclassified taxonomy (Fig 1a and Table S1). Approximately 88.9% of the bacterial genomes were derived from phyla of *Proteobacteria, Firmicutes,* and *Actinobacteria,* indicating that the WGD might be biased by the over sequencing of some specific taxa, especially those taxa of medical importance. To avoid such bias, individual genomes of the same species were merged together, resulting in a curated percentage of 23.6% bacterial species obtained from these three dominant phyla. Still, some genera displayed high prevalence of pathogenic species, such as *Klebsiella* (57.1%), *Enterobacter* (55.6%), and *Escherichia* (25.0%), compared to the average 8.6% of pathogenic species within *Bacteria.*

**Fig. 1.**
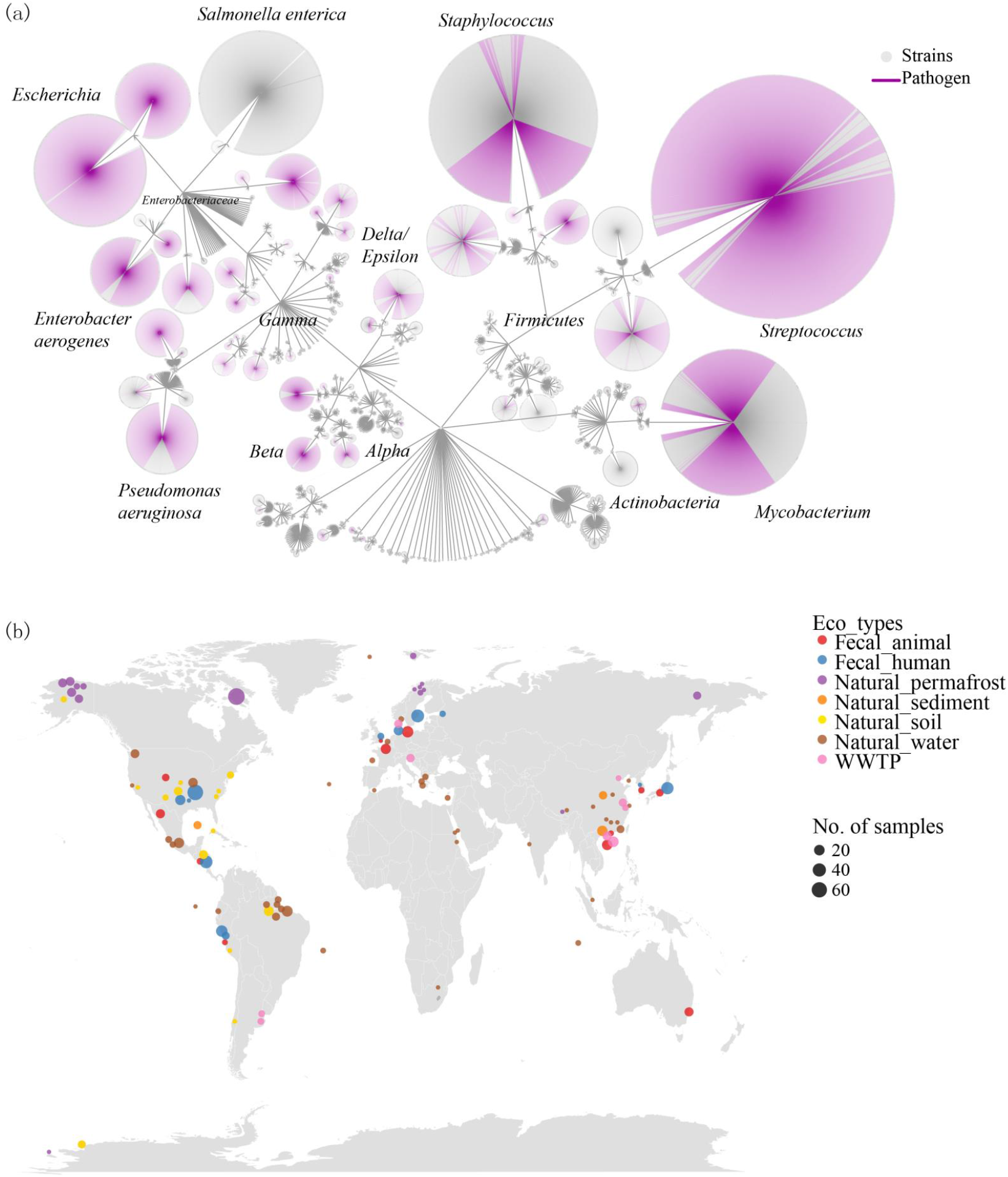
Overview of the Whole Genome Database (WGD) and Metagenome Database (MGD). (a) The phylogenetic relationship of all 54,718 bacterial genomes of 45 phyla in WGD and the occurrence of ARGs (blue nodes) and Rank I ARGs (red nodes). The pathogenic strains were indicated by purple edges. (b) The global map of metagenomic datasets in MGD. The size of the datasets (nodes) was proportional to the number of samples and the color was differentiated by the eco-type.

The MGD collected in this study covered a wild range and diversity of both the anthropogenic (WWTPs, animal feces and human feces) and natural (water, sediment, soil, and permafrost) habitats (Fig 1b and Table S2). The classification of the ecosystems was manually conducted based on a two-tier hierarchical classification system[61]. For a more comprehensive analysis and specific detailed comparison, each habitat (eco-type) was further classified into 3-5 eco-subtypes, except for the natural permafrost habitat category. The datasets in MGD are both geographically and ecologically distinct, collecting across different countries and continents, together to draw a global map.

Rarefaction curves [57] are adopted to evaluate the representativeness and coverage of these two datasets about the ARGs in the bacterial life tree and in the environments. The rarefaction curve for WGD was plotted by the consecutive addition of one random genome extracted from the collection of 54,718 bacterial genomes. The number of unique ARGs (not detected in those genomes previously added into the pool) provided by the new genome was counted. After the inclusion of the last genome, the rarefaction curve for WGD gradually reached a plateau of 2,625 unique ARGs (Fig S1a). Additional inclusion of new bacterial genomes in the future was expected to contribute a mild increase of 1 novel ARG per 170 genomes (minimum number of new genomes), indicating the representativeness of the current collection of bacterial whole genomes.

Similarly, for MGD, all the raw reads from 854 samples were pooled together to construct a rarefaction curve. The raw reads were randomly selected from the entire MGD pool one by one, to be evaluated against the SARG (as described in the section of methods). The total number of unique ARGs was added by one if the reference ARG assigned to this raw read was not identified previously. It seemed that after sampling 1.6E+10 raw reads from the entire MGD pool, the rarefaction curve illustrated a trend of a flat slope, and finally reached a plateau of 3,821 unique ARGs with the inclusion of all 2.4E+10 raw reads. It was predicted that one novel ARG could be expected with an extra sample of at least 1.4E+8 raw reads, also demonstrating the representativeness of the current collection of metagenomic datasets. Besides, the coverage of the MGD to the ARGs harbored by individual habitats was specifically evaluated by drawing the rarefaction curves for each eco-type. Generally speaking, the rarefaction curves for the anthropogenic habitats of animal feces, human feces and WWTPs tend to have reached their plateaus of 2,951, 2,815, and 3,131 unique ARGs (Fig S2), respectively. In contrast, the rarefaction curves were still growing gently for the natural environments of water (2,514 ARGs), permafrost (1,897 ARGs), soil (1,618 ARGs), and sediment (1,181 ARGs) after fully sampling all the raw reads. However, when calculating the new raw reads required to obtain one novel ARG, it was suggested that the current MGD had a higher representativeness to the habitats of WWTP (8.7E+7), human feces (2.4E+7), natural sediment (2.1E+7) than the habitats of animal feces (8.2E+6), natural water (4.2E+6), natural permafrost (3.8E+6), and natural soil (1.2E+6). The disagreement between two observations could be caused by the different patterns of the richness and density of the antibiotic resistome within 7 habitats. For example, the first inclusion of 1.0E+8 raw reads contributed to an average increase of 1,182 unique ARGs in the anthropogenic environments, which almost double the results (601 unique ARGs) of the natural environments. Another example was that at the level of 7.9E+8 raw reads (the lowest sampling depth), the anthropogenic environments recovered 2,250 unique ARGs while only 1,421 unique ARGs were detected in the natural environments. These two observations indicated the higher richness and higher density of antibiotic resistance in those anthropogenic habitats.

Overall, both current versions of WGD and MGD were evaluated to have high representativeness of the antibiotic resistance in the bacterial life tree and in the environments. However, additional sampling is expected to enrich the ARG profiles in individual natural habitats, especially the categories of water, permafrost, and soil.

### ARGs-OSP (antibiotic resistant genes-online searching platform)

The ARGs-OSP provides the search and download functionality of the occurrence and abundance of the ARGs and *intI1* genes retrieved by a global investigation in this study. The mothertables were separated into two Modules for the WGD and MGD, where the information of the hosts and the habitats can be specifically offered. In order to facilitate the diverse requirements of future studies, the identification and quantification of the target genes can be constrained by the customized cutoff selected by the users, including the identity, hit-length or hit-ratio, and e-value. Without any input, the ARGs-OSP returns the results of all target genes detected in all whole genomes or metagenomes, applying the default search cutoff described in the section of the methods. In each Module, three classes of inquiry factors are listed at the top middle panel, including the ARGs (*“Sequence “, “Subtype “* and *“Type”*), the host information (*“Genome”*, *“Accession”*, *“Organism”*, *“Assembly_level”*, *“Phylum”, “Class”, “Order”, “Family”, “Genus”, “Species”, “Strain”,* and *“Pathogen”*) for Module 1, the habitat information (*“Accession “, “Eco-subtype “* and *“Eco-type”*) for Module 2, and the searching criteria (*“Identity”, “Hit-length”* or *“Hit-ratio”* and *“E-value”*).

In Module 1 (Fig 2), the ARGs can be viewed and searched in all the whole genomes under a given cutoff, and all the details about their host taxa were summarized based on the annotation in GenBank and NCBI genome database. This functionality would be extremely beneficial for those research interests into the occurrence of ARGs in some specific bacterial lineages or the phylogenetic distribution of some specific ARGs. To meet versatile demands at the user end, the host information supports the details of the taxonomy information (from phylum level to strain level), the accession number (*“Genome”* for genbank assembly accession and *“Accession”* genbank sequence accession number), the status of the genomes *(“Assembly_level”* referring to the genome completeness), and the potential pathogenicity (*“Pathogen”),* which can all be set to filter the search results simultaneously. The searching criteria in Module 1 were transformed into aa-based *“Identity”* and *“Hit-ratio”* against the reference SARG, and the *“Hit-ratio”* referred to the percentage of hit-length to the reference length. Without imputing any searching criteria, ARGs-OSP will output all the inquiries that meet the default cutoff (≥ 90% aa identity, ≥80% aa hit-ratio and ≤ e-value 1e-5). The allowance of searching criteria is defined in the range of 50%-100% aa identity, 50%-100% aa hit-ratio and e-value smaller than 1.0E-1. For more comprehensive downstream analysis by the user end, the complete output table can be easily downloaded as a local file.

**Fig. 2.**
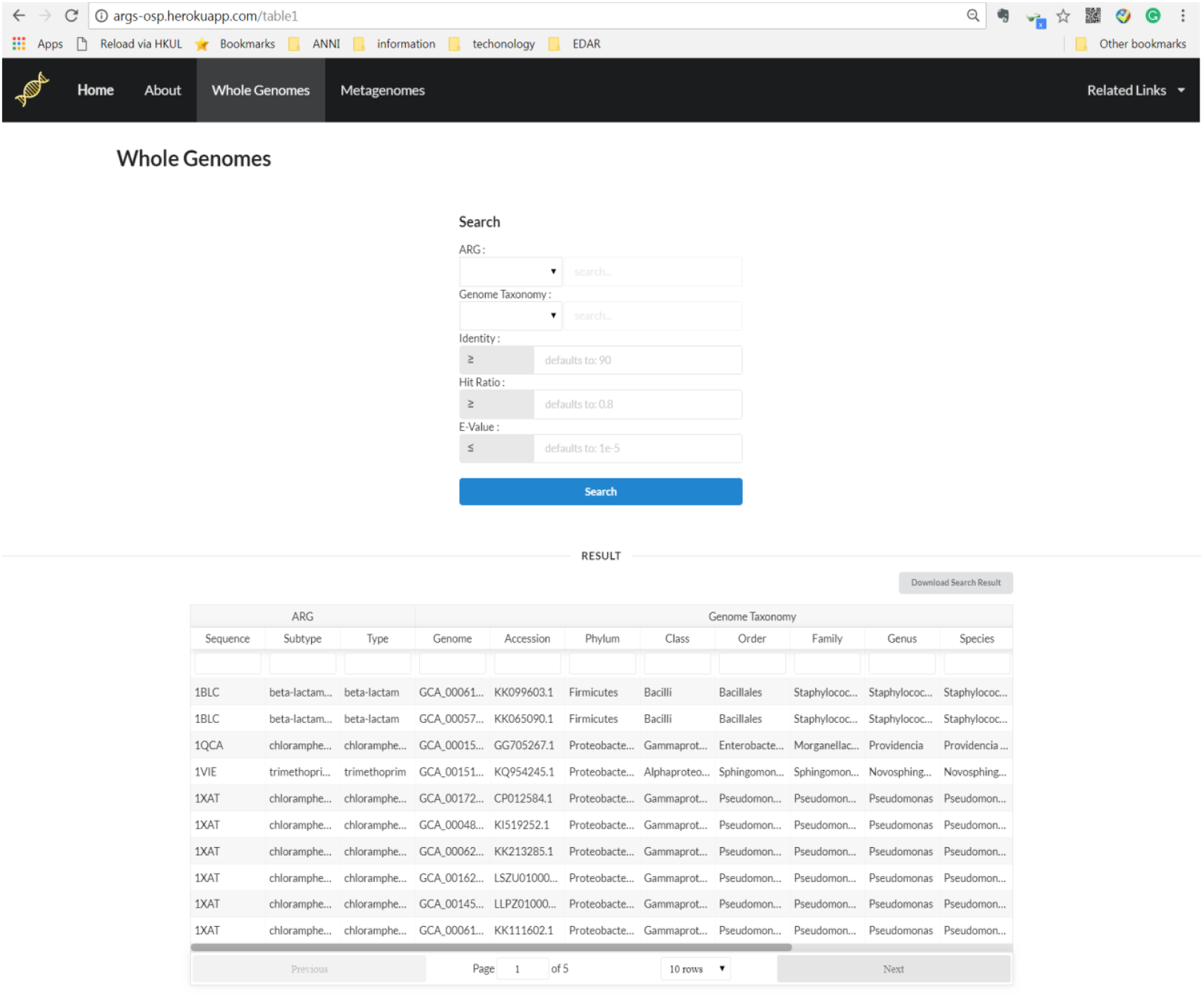
The layout of the ARGs-OSP Whole Genomes.

Module 2 (Fig 3) was constructed by investigating the ARGs in the collection of metagenomes under different combinations of cutoff, combined with the habitat information of each metagenomic dataset. This functionality would be extremely helpful for users inquiring the abundance of some specific ARGs in all the environments or the habitats of their interests, or for users to compare the ARG profile of local samples to a global collection. For reliable parallel comparison, users are recommended to process the ARG identification and annotation using the ARGs-OAP v1.0 [38, 45], which is compatible to the ARGs-OSP. In details, the habitat information covered the *“Accession”* (run accession number of NCBI SRA or MG-RAST databases), *“Eco-subtype “* and *“Eco-type”*, which was classified and curated manually. The searching criteria in Module 2 were also based on the aa *“Identity”* and *“Hit-length*” against the reference databases. The default criteria was set as ≥ 80% aa identity, ≥75% aa hit-ratio and ≤ e-value 1e-7, for empty cutoff input. The cutoff range allowed for searching is defined for the aa identity (60%, 70%, 80%, 90%, and 100%), aa hit-length (50%, 75%, and 100%) and e-value (1.0E-6, 1.0E-7, 1.0E-8, 1. 0E-9). However, compared to the identity, the hit-length and e-value were evaluated to have little influence on the ARG profile [45]. Since all the metagenomic datasets were trimmed into a standard read length of 100bp, the aa hit-length of 50%, 75% and 100% was expressed in the form of 17aa, 25aa and 33aa. As mentioned before, three abundance units (copy per cell, copy per 16S and ppm)[45] are provided for the flexible comparison at the user end, which could be easily switched by the buttons at the top left panel of output mothertable. A bottom panel is designed to customize the table layout and turn the pages.

**Fig. 3.**
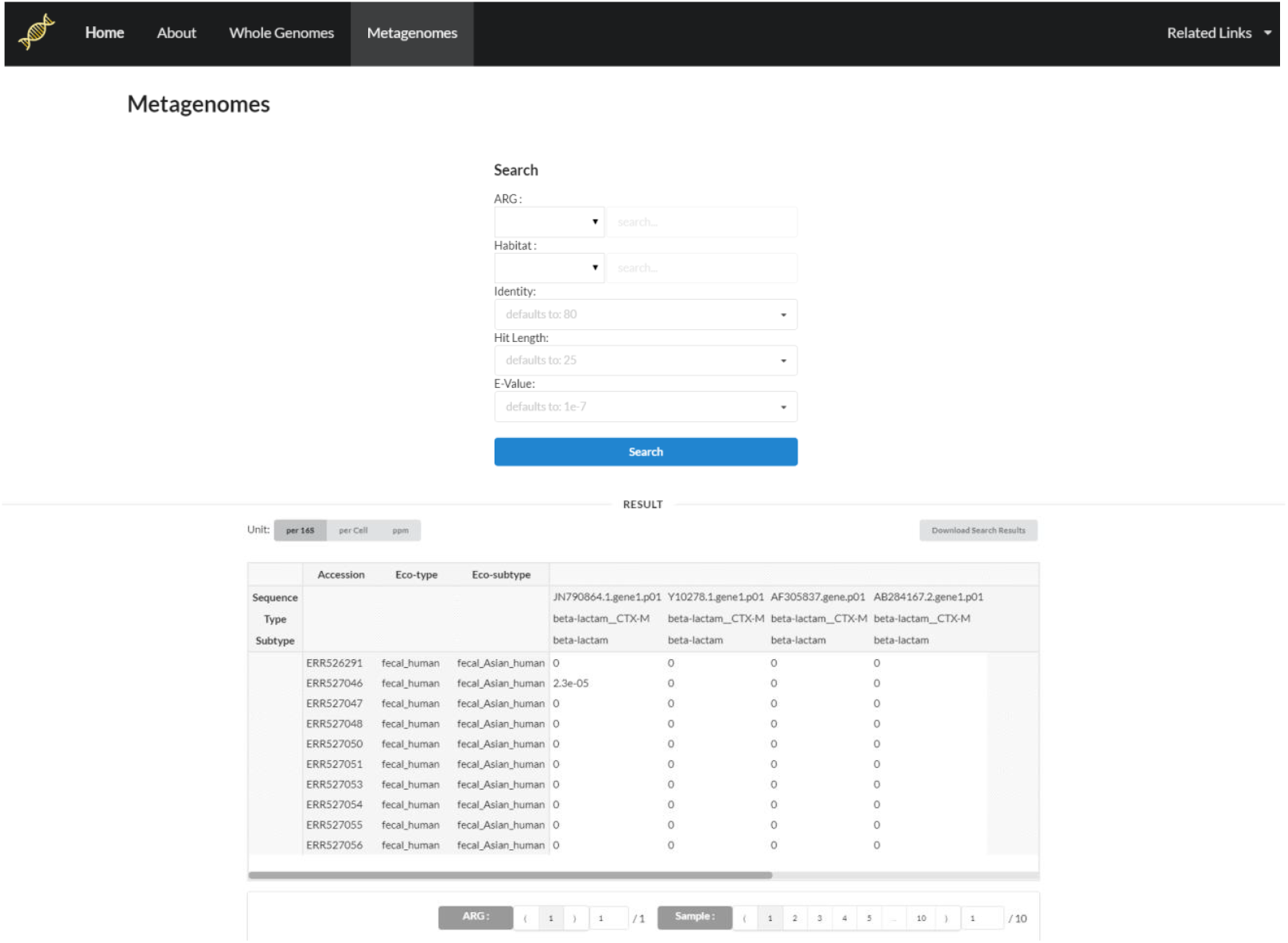
The layout of the ARGs-OSP Metagenomes.

### A global profile of antibiotic resistome

In this study, a global profile of antibiotic resistome was constructed and presented by integrating the phylogenetic and ecological distribution of ARGs from both WGD and WGD. In WGD, 2,625 ARGs (764 genotypes and 22 phenotypes) were detected in 809 bacterial species from 13 phyla. Within all the ARG-carrying species, 26.8% were identified as pathogenic species, which were almost 3 times the prevalence of pathogenic species in the WGD. This observation indicated that ARGs could be selected by the ecological fittings provided with strong selection force of anthropogenic pollution [62–66], where human pathogens are universally present. Among all the phenotypes, the multidrug (the occurrence of 29.4%), beta-lactam (15.5%), aminoglycoside (10.5%) and tetracycline (8.7%) were universally detected in bacterial species, which was consistent with a previous study investigating 2,500 complete genomes [47]. The ARG genotypes of *tetA, tetM, acrB, aph(3’)-I, aadA, mdtK*, *TolC*, and class A beta-lactamase resistant to tetracycline, aminoglycoside, multidrug and beta-lactam were frequently identified in 1.5% of bacterial species, covering a wide spectrum of taxonomy lineage and possessing more than 50% prevalence in human pathogens. These ARGs have successfully invaded across the phylogenetic barrier of the bacterial phylum level, especially into human microbiome, which should raise substantial alarm in both the medical and environmental fields.

3,821 ARGs from 993 subtypes/genotypes and 24 types/phenotypes were identified in diverse environments of the MGD in total. Even though the anthropogenic environments were illustrated to have ARGs of higher density and richness than the natural environments (Fig S2), after normalizing against the bacterial cell number in each sample, the abundance of total ARGs showed no significant difference (less than 10 folds) among the 7 eco-types (Fig S3). This observation suggested that those divergences of the density and richness of antibiotic resistant profiles could be caused by the density and richness of the microbial community among different eco-types, while the resistant level within the bacterial cells seemed to be quite stable.

Generally speaking, ARGs are widely distributed in almost all natural and anthropogenic habitats with the average abundance varying from 5.1E-2 copy per cell to 6.7E-1 copy per cell. It was not surprising since antibiotic resistance is originally harbored by natural microorganisms and most antibiotics used nowadays are produced by natural antibiotic producers [19]. The animal feces harbored the highest abundance of ARGs (6.7E-1 copy per cell) under the strong selective pressure of over-dosing antibiotics [67–69], which may promote the dissemination of ARGs [70]. The other two anthropogenic environments, the human feces and WWTPs, displayed similar level of ARGs of 3.7E-1 and 1.7E-1 copy per cell. It was notable that high abundance of ARGs was hosted by the microbial communities in the natural environments of permafrost (6.1E-1 copy per cell) and soil (2.6E-1 copy per cell), even denser than the WWTPs as the hotspot of ARGs [13, 22]. Within the soil eco-type, the eco-subtype of rural soil could be contaminated by the agricultural usage of animal manure, the other three soil eco-subtypes of city, amazon catchment and prairie displayed equivalent level of antibiotic resistant. A relative low level of ARGs was detected in the natural water (1.0E-1 copy per cell) and sediment (6.7E-2 copy per cell), while two obvious trends of heterogeneous and homogeneous distribution were respectively observed in the water and soil eco-types.

To make the results comparable to previous studies, the abundance of ARGs was transformed into the units of copy per 16S and ppm, and provided on the ARGs-OSP. Most habitats exhibited similar resistance level compared to previous studies [7, 54, 71] except for a much higher detection of ARGs in soil (2.6E-1 copy per 16S) and surface water (1.0E-1 copy per 16S) compared with the abundances of 2.0E-2 to 9.0E-3 copy per 16S and 2.0E-2 to 8.0E-3 copy per 16S, respectively. This difference could be contributed by the heterogeneity within the soil environments and divergence within the surface water environments, and a large sample size with multiple eco-subtypes in this study would help construct a more comprehensive and representative global profile.

Besides the abundance of total ARGs, the antibiotic resistome of different habitats also displayed their divergence regarding to their composition and diversity. Instead of directly counting the unique reference sequences detected in one eco-type, ARGs conducting the same resistant mechanisms were classified into one genotype. The sediment environment harbored a deficient collection of 281 ARG genotypes, accounting for less than 30% of the global profile. Approximately 50% of the global resistant profile was recovered in the soil (404 genotypes), permafrost (513 genotypes) and human feces (604 genotypes). The animal feces, water and WWTP environments provided a rich pool of antibiotic resistance, covering 714, 722 and 761 genotypes, respectively. Those genotypes that have long been the focus of ARG-related researches were found to be widespread and abundant in all 7 eco-types, including aminoglycoside resistance gene *aph(3)-I,* the beta-lactam resistant gene *TEM*, sulfonamide resistance genes of *sul1* and *sul2,* MLS resistant genes of *macA, macB,* tetracycline resistant genes of *tetA* and *tetM,* multidrug resistant genes of *acrA* and *acrB,* and vancomycin resistant gene *vanA* [14, 54, 72]. Overall, the anthropogenic pollution appeared to have a weak influence on the total antibiotic resistant level within the bacterial cell, but may cast a relatively strong impact on the diversity and density of the antibiotic resistome within the habitat.

### Co-occurrence of the ARGs and intI1 genes: a demonstration of ARGs-OSP

The *intI1* genes were considered as a potential indicator for anthropogenic pollution [73] because of its high abundance and universal occurrence in human-related environments, such as wastewater treatment plants and animal feces [74, 75]. Previous studies on the correlation between *intI1* genes and human pollution mainly focused on three directions: 1) *intI1* genes tended to have high abundance in human-related environments, and cannot be effectively removed from WWTPs [76–78]; 2) the co-selection of *intI1* genes with ARGs, metal resistant genes (MRGs) and disinfectant resistant genes [79–81]; 3) the abundance of *intI1* genes increases with anthropogenic pollution, such as heavy metal, disinfectants, antibiotics, and pesticides [82–84]. Besides, some ARGs *(sul1, sul2* and *tetM)* were evaluated to have strong and positive correlation with *intI1* genes in polluted sediment samples [85]. However, the co-occurrence of the *intI1* genes and ARGs was not comprehensively evaluated before. Moreover, this co-occurrence could be casually caused by their co-selection by the same selective pressure or by physically linked on the same transposons and plasmids, and thus resulted in quite inconsistent relationship.

To evaluate whether the *intI1* genes could be an indicator for the anthropogenic pollutant of total ARGs and in which habitats there could be a strong correlation, the mothertables of ARGs and *intI1* genes were downloaded from ARGs-OSP. The abundances of the total ARGs and total *intI1* genes were summed up for each sample, and the samples were pooled into 7 eco-types. The second question was raised in this study that if the *intI1* genes were a weak proxy for total ARGs, which subgroup of ARGs could be indicated by the *intI1* genes? One subgroup of 107 ARGs found on class 1 integrons in WGD (manuscript under review) was proposed as a potential target.

Overall, the correlation of *intI1* genes to two groups of ARGs (total ARGs and ARGs on class 1 integrons) in 7 habitats showed that higher abundance of ARGs corresponding to higher abundance of *intI1* genes (Table 1). However, it was interesting to find that high abundance of total ARGs was detected in many samples, while no *intI1* gene was identified (light blue nodes in Fig 4). This indicated that *intI1* genes were not a universal marker for the presence of ARGs, not even for the anthropogenic environments where *intI1* gene was absent in a large portion of animal fecal and human fecal samples. This observation was also supported by the co-occurrence of *intI1* genes and ARGs in the WGD, where *intI1* genes were highly conserved in the class of *Gammaproteobacteria* (red nodes in Fig 5). Nevertheless, the ARGs were discovered to be widely distributed across different classes (light blue nodes in Fig 5), and the antibiotic resistance carried by those bacterial species outside the spectrum of *Gammaproteobacteria* were not likely to be indicated by the *intI1* genes.

**Fig. 4.**
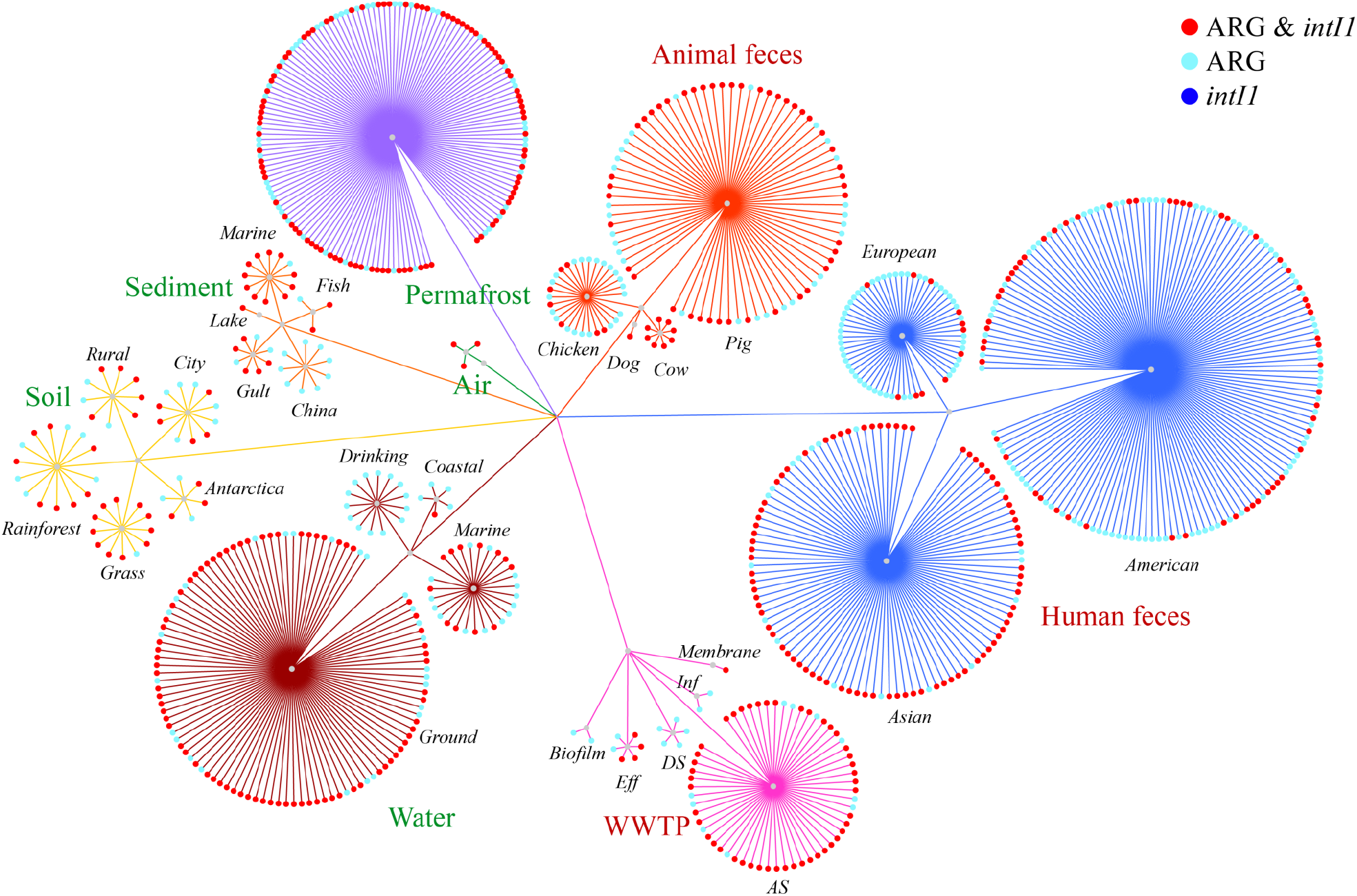
The ecological co-occurrence of class 1 integrases and total ARGs in 7 eco-types.

**Fig. 5.**
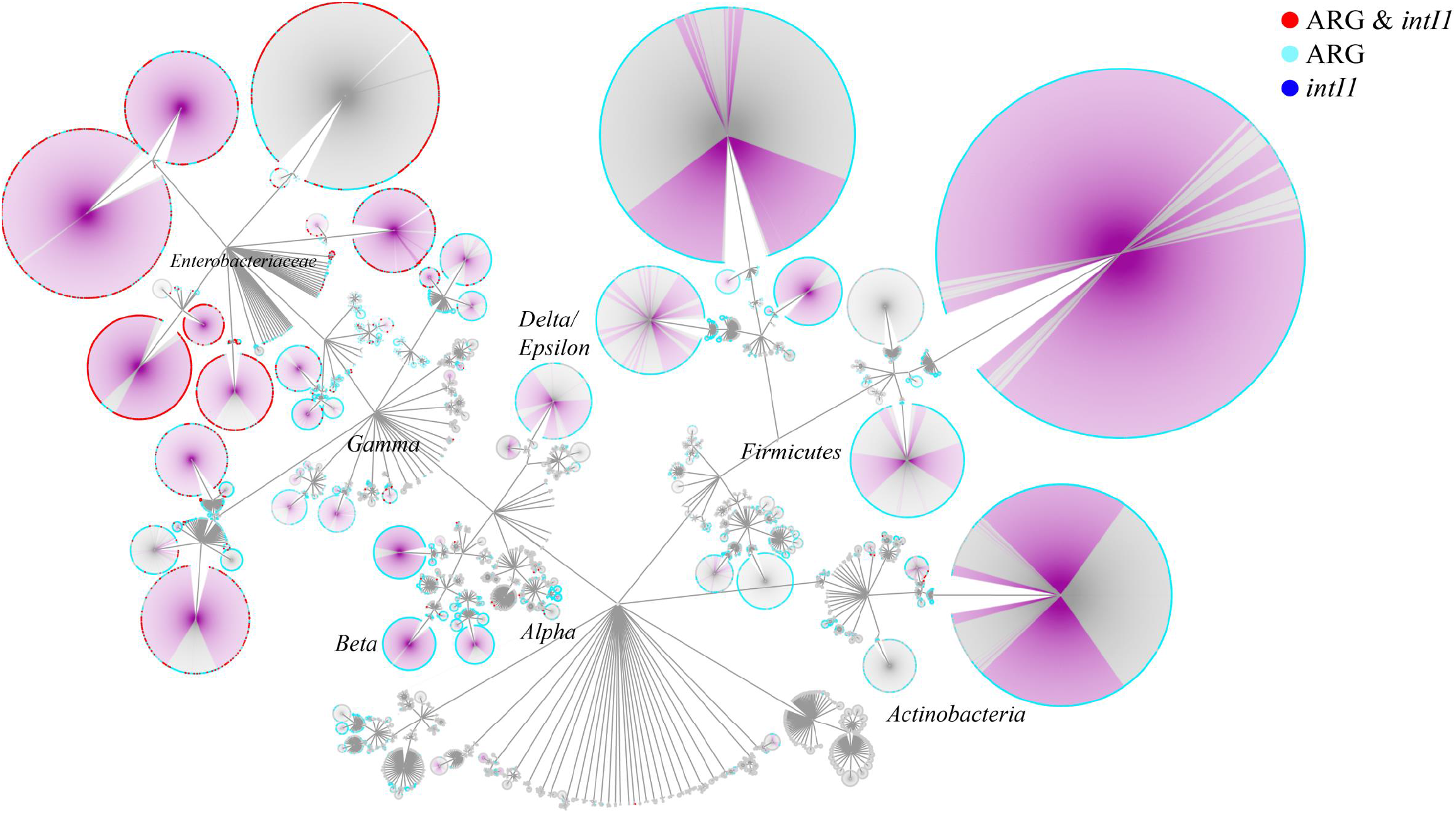
The phylogenetic co-occurrence of class 1 integrases and total ARGs in bacterial life tree.

**Table 1.** The linear regression relations of the abundance of *intI1* genes to the abundance of total ARGs (Total) and 107 ARGs detected on the class 1 integrons (Class I).

| Eco-type | R<sup>2</sup> |  | Slope |  |
| --- | --- | --- | --- | --- |
|  | *intII*-Total | *intII*-Class I | *intII*-Total | *intII*-Class I |
| Natural_Permafrost (144) | 0.08 | 0.09 | 0.69 | 0.28 |
| Natural_Sediment (33) | 0.37 | 0.21 | 2.00 | 0.39 |
| Natural_Soil (52) | 0.06 | 0.44 | 0.53 | 0.61 |
| Natural_Water (146) | 0.34 | 0.35 | 0.83 | 0.64 |
| WWTP (69) | 0.03 | 0.22 | 0.62 | 0.95 |
| Fecal_Animal (110) | 0.47 | 0.75 | 2.10 | 1.10 |
| Fecal_Human (300) | 0.30 | 0.35 | 1.40 | 0.63 |
| All (854) | 0.04 | 0.35 | 0.40 | 0.65 |

Besides an overall picturing of their correlation, the linear regressions were fitted to the *intI1* genes and total ARGs in 7 eco-types, which displayed week linear relation with the R varied from 0.03-0.47 (Table S1). Here, to avoid the biased caused by sequencing depth, samples with no presence of *intI1* gene were not considered during assessment. The low R^2^ of WWTP (0.03), soil (0.06) and permafrost (0.08), indicated that the *intI1* genes had a poor linear relationship to the total ARGs in these habitats and their role as an indicator should be treated with caution. Moreover, 7 habitats were found to comply with different linear relations, regarding to their slopes of 0.53 to 2.10. The anthropogenic environments were expected to have higher slope of linear relations because of higher level of anthropogenic pollution, while this trend was not clear in this study. Thus, the correlation between *intI1* and the total ARGs was both weak (in terms of R^2^) and inconsistent (in terms of slopes) for different environments.

The *intI1* genes were fit with better linear relationships (in terms of R^2^) to the subgroup of ARGs on class 1 integrons in all habitats except for the sediment (Table 1 and Fig S4). The *intI1* genes could be proposed as a good indicator for ARGs on class 1 integrons, especially for animal feces (R of 0.75). Even though, this relationship may only be applied to samples in soil, water, human feces and animal feces (R^2^ ≥ 0. 35), and the linear relationship should also be specifically tuned for each habitat.

## Conclusions

In this study, a global profile of the antibiotic resistome was constructed by integrating two big datasets of the WGD (54,718 bacterial genomes) and MGD (854 metagenomes of 7 habitats). Both the WGD and MGD were evaluated to have good representativeness and comprehensive coverage of ARGs in bacterial genomes and metagenomes, serving as the fundamental bases to investigate the phylogenetic and ecological distribution of antibiotic resistance. Moreover, all ARGs were identified and quantified using a standardized pipeline for reliable parallel comparison. Most importantly, a user-friendly and well-organized online searching platform, the ARGs-OSP, was designed to publish all the mothertables obtained in this study, making the data easily accessible for other researchers. The ARGs-OSP can serves as valuable sources and references for future studies with versatile research interests, while avoiding unnecessary re-computations. Finally, the potential of the ARGs-OSP was demonstrated by evaluating whether the *intI1* genes could be a good proxy for the anthropogenic pollution of ARGs by investigating their co-occurrence.

The major limitation of this study lies in the two datasets of WGD and MGD. The WGD could be biased by over-sequencing those bacterial species of research interest and medical importance, which may not represent all the environmental microbes. For the MGD, the current sampling scheme mainly focused on the representative eco-types, and the extreme environments were not included in this version. Besides, the availability of qualified samples was limited for some important environments, such as air. However, with the rapid development and decreasing cost of sequencing techniques, datasets of high quality and massive quantity is promising for the expansion of WGD and MGD. Further updates of ARGs-OSP is expected to be enhanced responding to the continuously growing number of bacterial genomes and environmental metagenomes in public database. Also, more flexible search and visualization functionality will be continuously complemented in future versions of ARGs-OSP, for more convenient and versatile usage.

## Declarations

### Availability of supporting data and materials

The datasets supporting the results of this article are available on the ARGs-OSP (XXXX)

### List of abbreviations

ARGs: antibiotic resistant genes
ARGs-OAP: ARGs online analysis pipeline
ARGs-OSP: ARGs online searching platform
*intI1*: class 1 integrases
WGD: whole genome database
MGD: metagenome database.

### Author’s Contributions

A. N. Zhang downloaded the datasets, analyzed the data, designed the platform and wrote the manuscript. Chen-Ju Hou constructed the platform. L. L. Guan provided suggestions in the data analysis and manuscript preparation. T. Zhang guided webpage development and revised this manuscript.

---

**Fig. S1.**
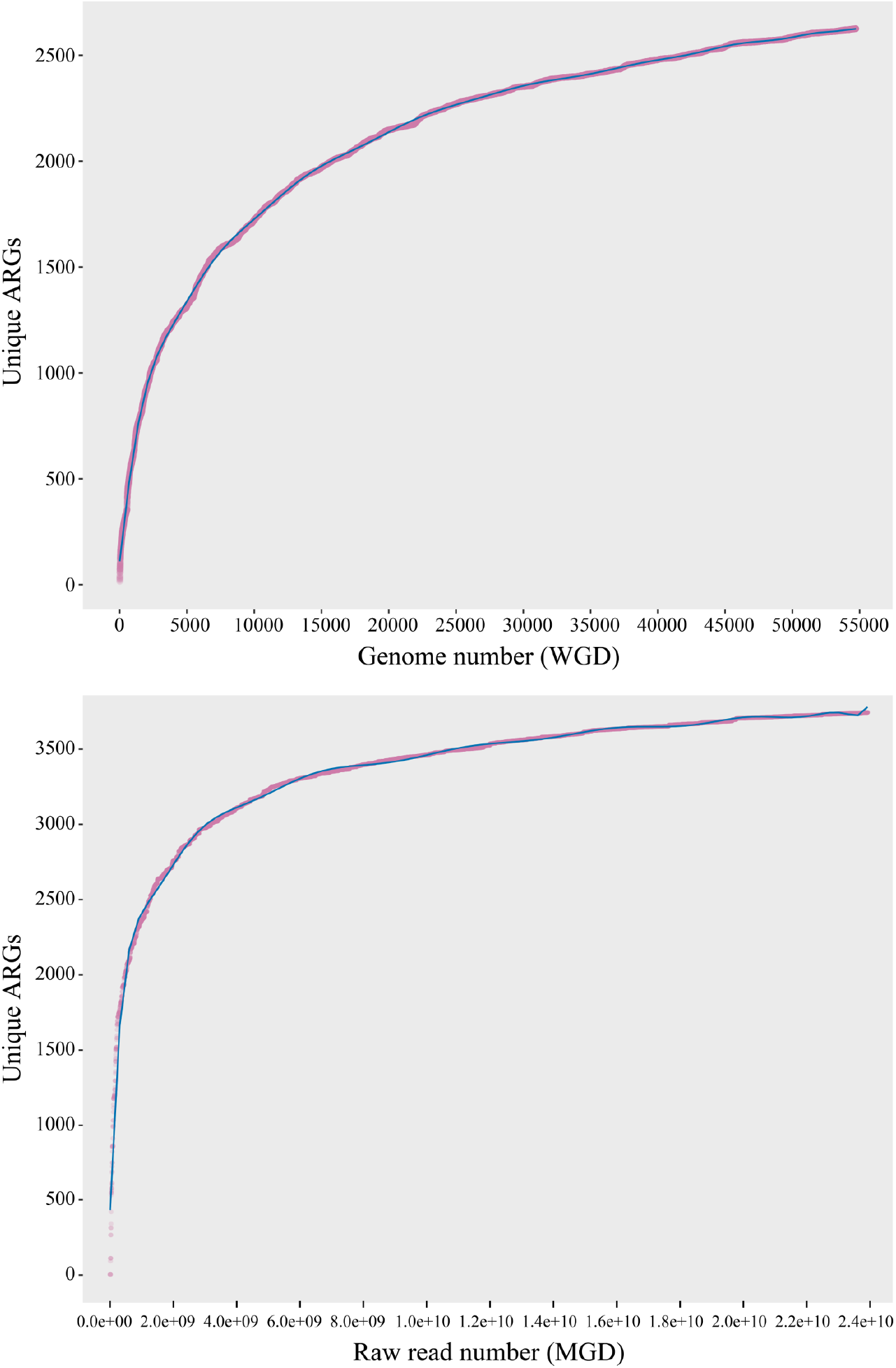
The rarefaction curves of the unique ARGs in the Whole Genome Database (WGD, each genome) and Metagenome Database (MGD, each raw read), constructed by randomly subsampling without replacement.

**Fig. S2.**
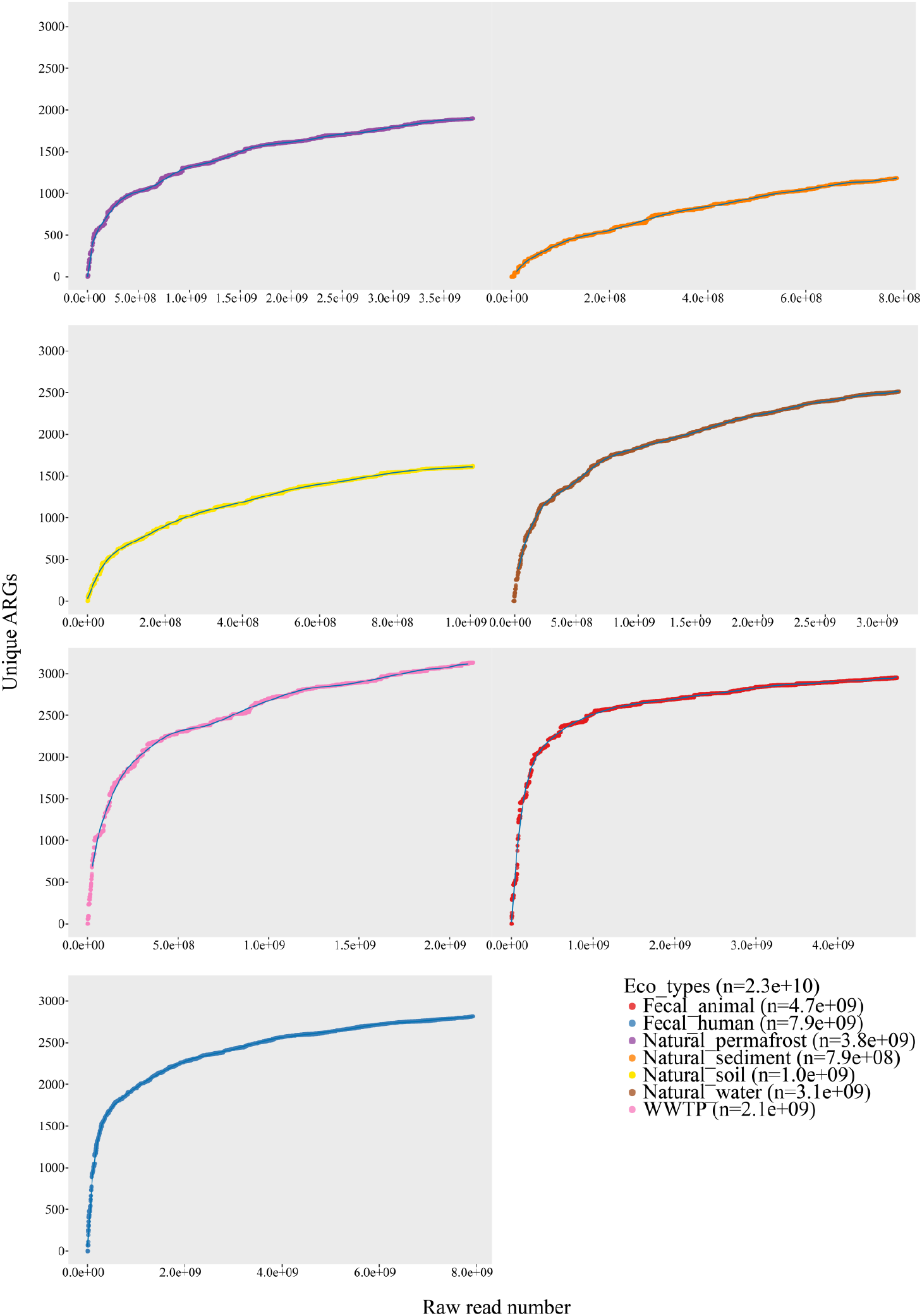
The rarefaction curves of the unique ARGs constructed specifically for 7 eco-types of Metagenome Database (MGD, each raw read) by randomly subsampling without replacement.

**Fig. S3.**
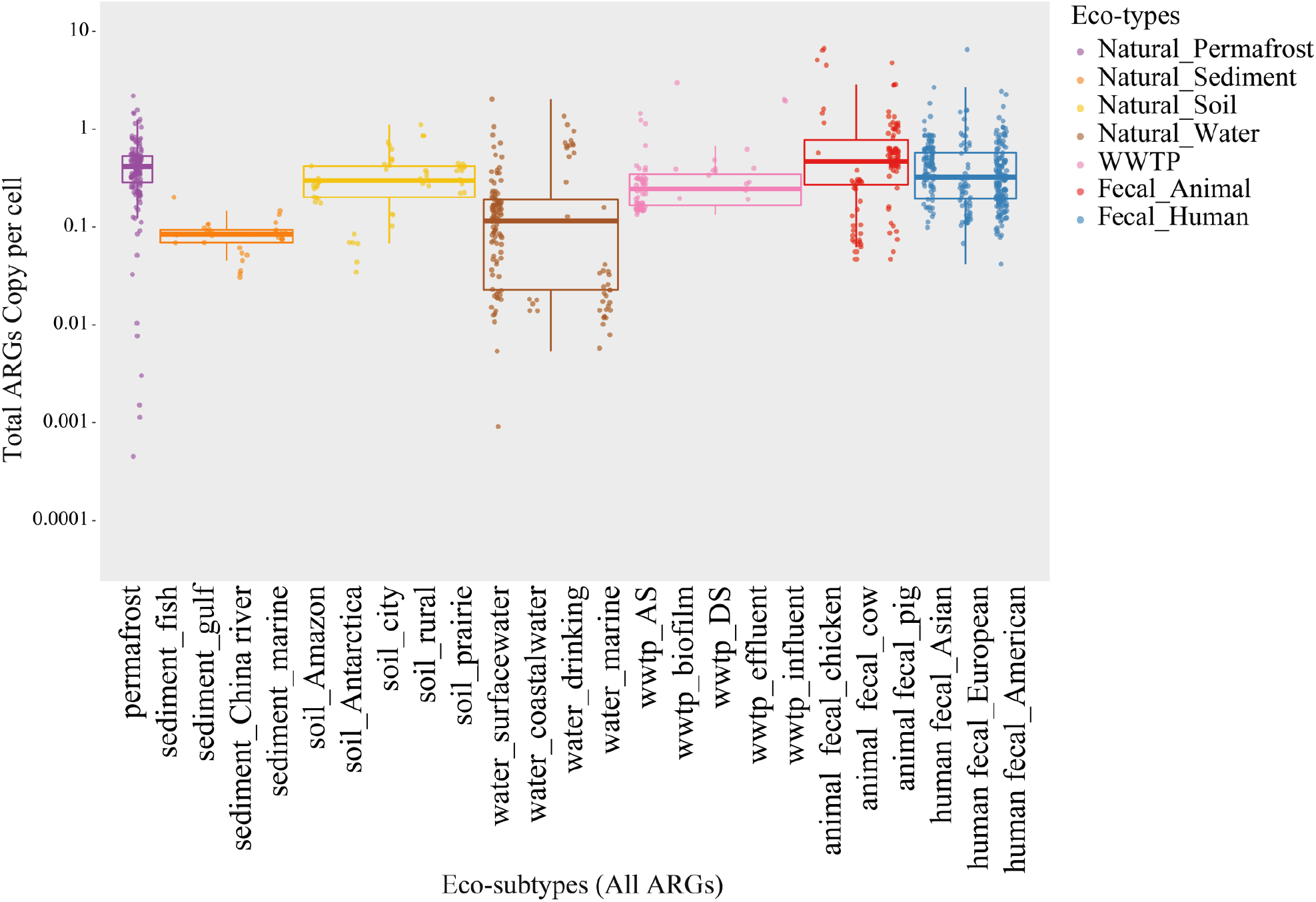
The ecological distribution and abundance (copy per cell) of total ARGs in 25 eco-subtypes.

**Fig. S4.**
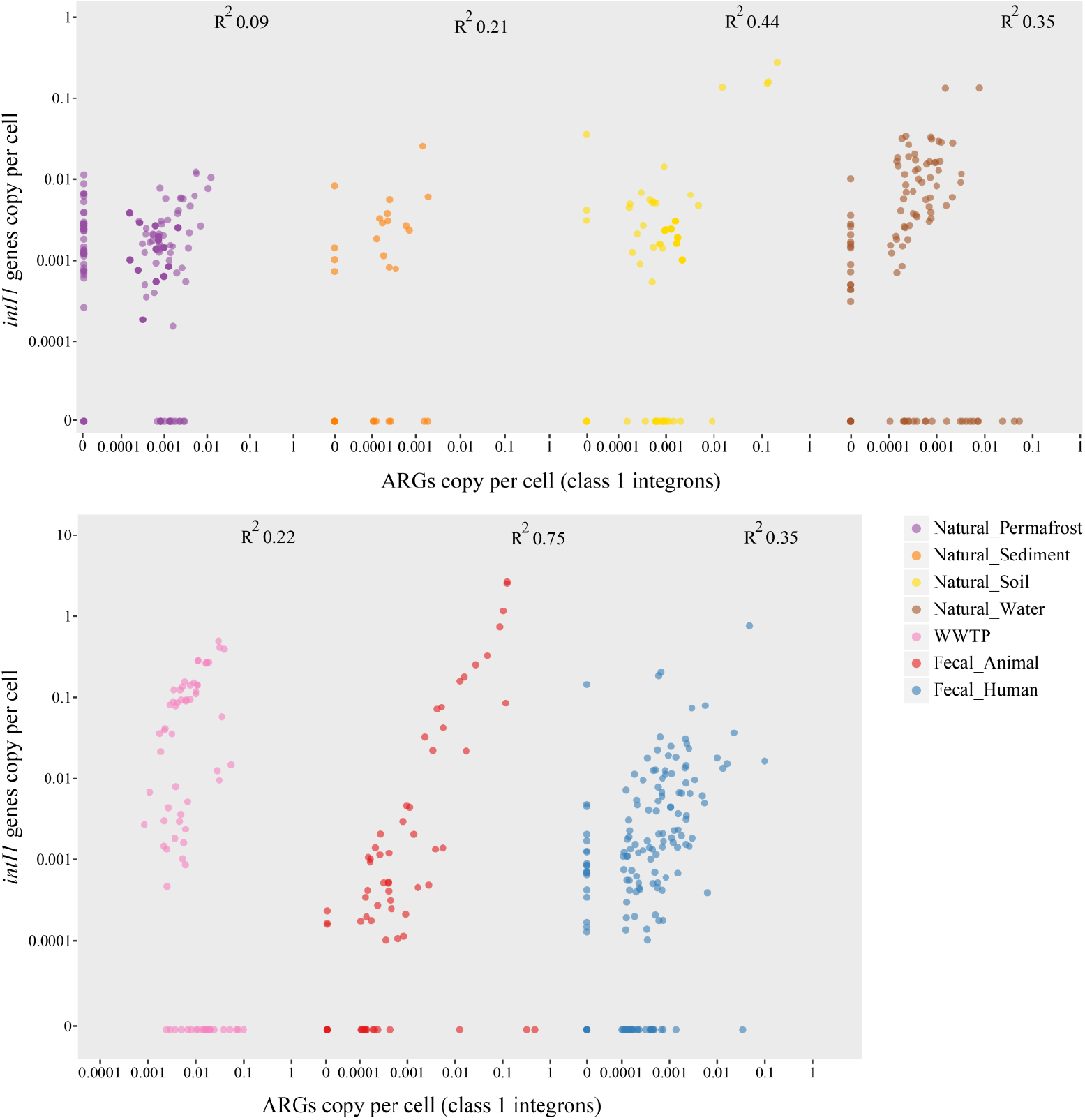
The co-occurrence of class 1 integrases and a subgroup of ARGs that were discovered on class 1 integrons in 7 eco-types, fitted by linear regression (R).

